# Negative regulation of *C. elegans* innate immunity by orphan nuclear receptor NHR-42

**DOI:** 10.1101/2022.09.12.507697

**Authors:** Debanjan Goswamy, Sid A. Labed, Xavier Gonzalez, Javier E. Irazoqui

## Abstract

Positive and negative regulators of innate immunity work together to maintain immune homeostasis. We previously discovered that HLH-30/TFEB is a critical transcription factor that positively regulates host defense genes upon *S. aureus* infection in *C. elegans*. However, repression of host defense genes and negative regulation of immunity remain poorly understood. In this study, we identified *nhr-42* as a negative regulator of host defense genes functioning downstream of HLH-30/TFEB, with major implications in host survival and metabolism after infection. *nhr-42* expression is induced in an HLH-30/TFEB dependent manner mostly in the pharynx upon infection. We find that animals lacking *nhr-42* have higher expression of host defense genes, which enables enhanced survival after infection. Antimicrobials expressed in the pharynx such as *abf-2*, function downstream of *nhr-42* to confer resistance to infection by mitigating pathogen burden. Furthermore, *nhr-42* deficient animals are defective in lipid mobilization, having higher lipid stores compared to wild type animals after infection. *nhr-42* therefore enables *C. elegans* to limit the host defense response and reallocate energy resources through lipid mobilization after infection. To our knowledge, this is the first report of a transcription factor that represses host defense genes in *C. elegans*.

## Introduction

Detection of pathogenic signals by an organism leads to extensive changes in cellular transcriptional programs mediated by transcription factors. In both invertebrate and vertebrate cells transcription factors induce host defense gene expression to combat the threat of pathogenic infection (Cohen and Troemel, 2015; Lemaitre and Hoffmann, 2007). Conversely, failure to downregulate inflammatory genes leads to autoinflammation and tissue damage, which are detrimental to the host (O’Shea et al., 2002). Negative regulators of innate immunity are crucial because they are vital to maintaining host immune homeostasis. Previous research has discovered several transcription factors that upregulate innate immune genes, metabolic genes, and other processes required to combat infection (O’Neill et al., 2016). However, our understanding of how positive and negative regulators of immunity work to differentially regulate genes to maintain immune homeostasis is limited.

*Caenorhabditis elegans* can mount pathogen-specific transcriptional host defense responses (Irazoqui et al., 2010), be monitored for metabolic and reproductive dysfunction, and be genetically manipulated to identify regulators of complex biological processes. In the past two decades, several evolutionarily conserved host defense pathways that induce host defense genes in *C. elegans* were identified (Martineau et al., 2021). In contrast, how host defense genes are repressed by negative regulators of innate immunity in *C. elegans* remains unknown.

Previously, we discovered that a *C. elegans* transcription factor called Helix-loop-helix-30 (HLH-30), which is orthologous to human Transcription Factor EB (TFEB), is essential for upregulation of host defense genes, lipid catabolism, and autophagy during infection and that TFEB performs similar functions in murine macrophages (Visvikis et al., 2014). During *Staphylococcus aureus* infection HLH-30 upregulates host defense genes such as lysozymes, antimicrobial peptides, and C-type lectins (Visvikis et al., 2014). Animals lacking *hlh-30* are severely defective in survival of infection. However, how much of the host response is directly regulated by HLH-30 is unknown. We previously found that ∼80% of *C. elegans* genes induced upon infection are *hlh-30* dependent, but just ∼10% are estimated to be direct targets (Visvikis et al., 2014).

Among the genes induced by *hlh-30* are seventeen transcription factors, which could be downstream mediators of HLH-30 regulated gene expression. Ten of these transcription factors are predicted nuclear hormone receptors (NHRs). *C. elegans* has an expanded family of 284 NHRs, compared to 48 in humans and 49 in mice, but only around 20 of these NHRs have a known function. NHRs have been implicated in metabolism, reproduction, lifespan, and development in *C. elegans* (Antebi, 2015). They can induce or repress transcription based on their structure, protein-protein interactions, and ligand binding (Hoffmann and Partridge, 2015). Recent studies have only begun to uncover the role of these transcription factors in host defense (Peterson et al., 2019).

In this study we provide evidence that *nhr-42* acts downstream of *hlh-30* to repress the host response to infection, with key roles in infection survival and lipid mobilization. To our knowledge, this is the first report of a negative regulator of host defense genes in *C. elegans*.

## Results

### *S. aureus* induces *nhr-42* in an *hlh-30* dependent manner

We previously showed that *hlh-30* induces around 600 genes during *S. aureus* infection (Visvikis et al., 2014). However, HLH-30 is predicted to induce just 10% of these genes by directly binding their promoter region (Visvikis et al., 2014) (Figure 1a). We hypothesized that indirect regulation of host defense genes by *hlh-30* could be mediated by any of the seventeen transcription factors it induces upon infection (Visvikis et al., 2014) (Figure 1a). The site of *S. aureus* infection in *C. elegans* is the intestine. To identify their biological significance, we knocked down each of the transcription factors by RNAi, specifically in the intestine. We found that *nhr-42* RNAi significantly boosted host survival (Supplemental Figure 1d). Previous studies suggested that HLH-30 may bind directly to the promoter region of *nhr-42* (*Unlocking the secrets of the genome*, 2009) and therefore may directly regulate its expression during infection (Figure 1b).

**Figure 1.**
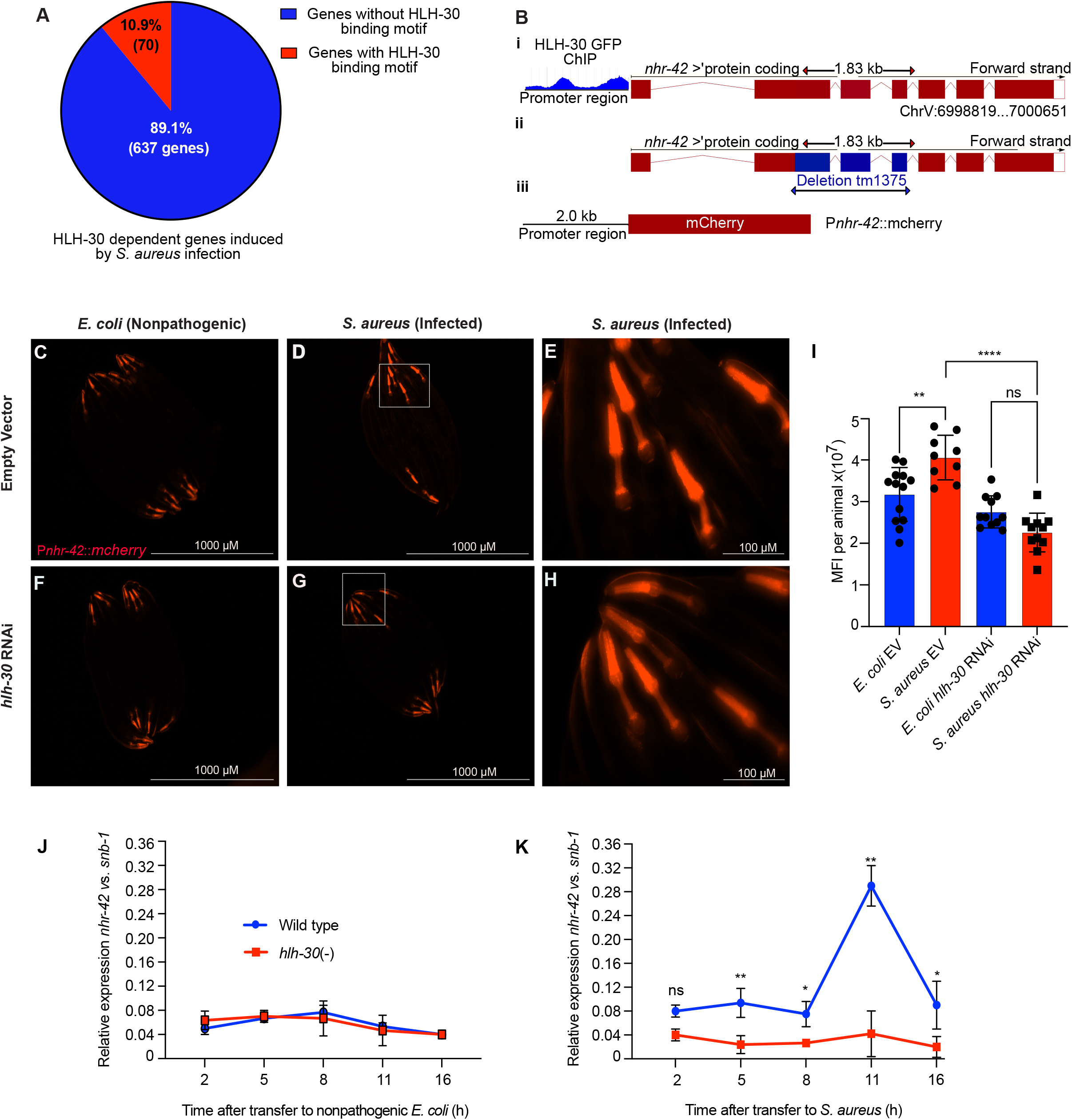
*S. aureus* induces *nhr-42* in an *hlh-30* dependent manner. **(A)** Venn diagram of direct and indirect induction of genes by HLH-30 upon *S. aureus* infection. Data generated from RNA-seq analysis(Visvikis et al., 2014). **(B)**NHR-42 gene structure, i) promoter region showing HLH-30 binding site. GFP-ChIP peaks within 2000bp of the transcription start site (*Unlocking the secrets of the genome*, 2009), ii) tm1375 deletion of 451 bp spanning exon 2-4, iii) P*nhr-42*:mcherry reporter construct. **(C-H)** Epifluorescence micrographs of P*nhr-42*::*mcherry* animals fed *E. coli* expressing RNAi empty vector or *hlh-30* double stranded DNA, and then exposed to *E. coli* OP50 (C,F) or to *S. aureus*. SH1000 (D,G) for 24 h. E and H show zoomed in portions pharynx/pharyngeal intestinal valve of D and G respectively. Scale bar = 1,000 micrometer for images C, D, F, G and 100 micrometers for E and H. Data are representative of 2 independent replicates. 10-20 animals each. **(I)** Quantification of mean fluorescence intensity of (A-D). ** *P* ≤ 0.01, **** *P* ≤ 0.0001. One-way ANOVA. **(J-K)** RT-qPCR of *nhr-42* transcript in wild type and *hlh-30* loss of function mutants fed *E. coli* OP50 (J) or infected with *S. aureus* (K). * *P* ≤ 0.05, \*\**P* ≤ 0.01. *t*-test. 2-4 biological replicates. n=∼1000 animals per replicate.

To better understand the expression timeline of *nhr-42* in infected and uninfected animals we quantified mRNA levels of *nhr-42* in wild type and *hlh-30* mutants. *nhr-42* expression remained unchanged in uninfected wild type and *hlh-30* mutants (Figure 1j). In contrast, *nhr-42* expression was two-fold higher in wild type animals compared to *hlh-30* mutants after 5 h, 8 h, 11 h, and 16 h infection (Figure 1k). The peak in expression of *nhr-42* was seen after 11 h of infection in wild type animals and this peak in expression was absent in *hlh-30* mutants (Figure 1k). To determine the expression pattern of *nhr-42* in uninfected and infected animals, we constructed a transcriptional reporter strain with the native *nhr-42* promoter driving the expression of mCherry, P*nhr-42::mCherry* (Figure 1b). In this strain, *nhr-42* was primarily expressed throughout the pharynx and the pharyngeal-intestinal valve (Figure 1c). We used the *nhr-42* reporter strain to confirm *hlh-30* dependent induction of *nhr-42* during *S. aureus* infection. We found that mCherry fluorescence was significantly higher during infection in animals fed RNAi bacteria expressing empty vector compared to animals fed *hlh-30* RNAi. (Figure 1d, g, and i). Together these data suggested that pharyngeal induction of *nhr-42* during infection is *hlh-30* dependent.

### *nhr-42* represses infection resistance

A previous study found that *nhr-42* is expressed in the pharynx, intestine, and hypodermis (Reece-Hoyes et al., 2007). To systematically identify the tissues in which *nhr-42* functions for host defense, we performed tissue-specific knockdown of *nhr-42* by RNAi. *nhr-42* depletion in the intestine and whole animal resulted in significantly enhanced survival of *S. aureus* infection compared to vector controls (Figure 2a, b).Hypodermal depletion of *nhr-42* caused mild resistance (Figure 2c) and depletion in the muscle had no significant effect (Figure 2d). We obtained similar results with an *nhr-42* deletion mutant (Figure 2e). These data suggested that *nhr-42* loss enables *C. elegans* to survive *S. aureus* infection better than wild type.

**Figure 2.**
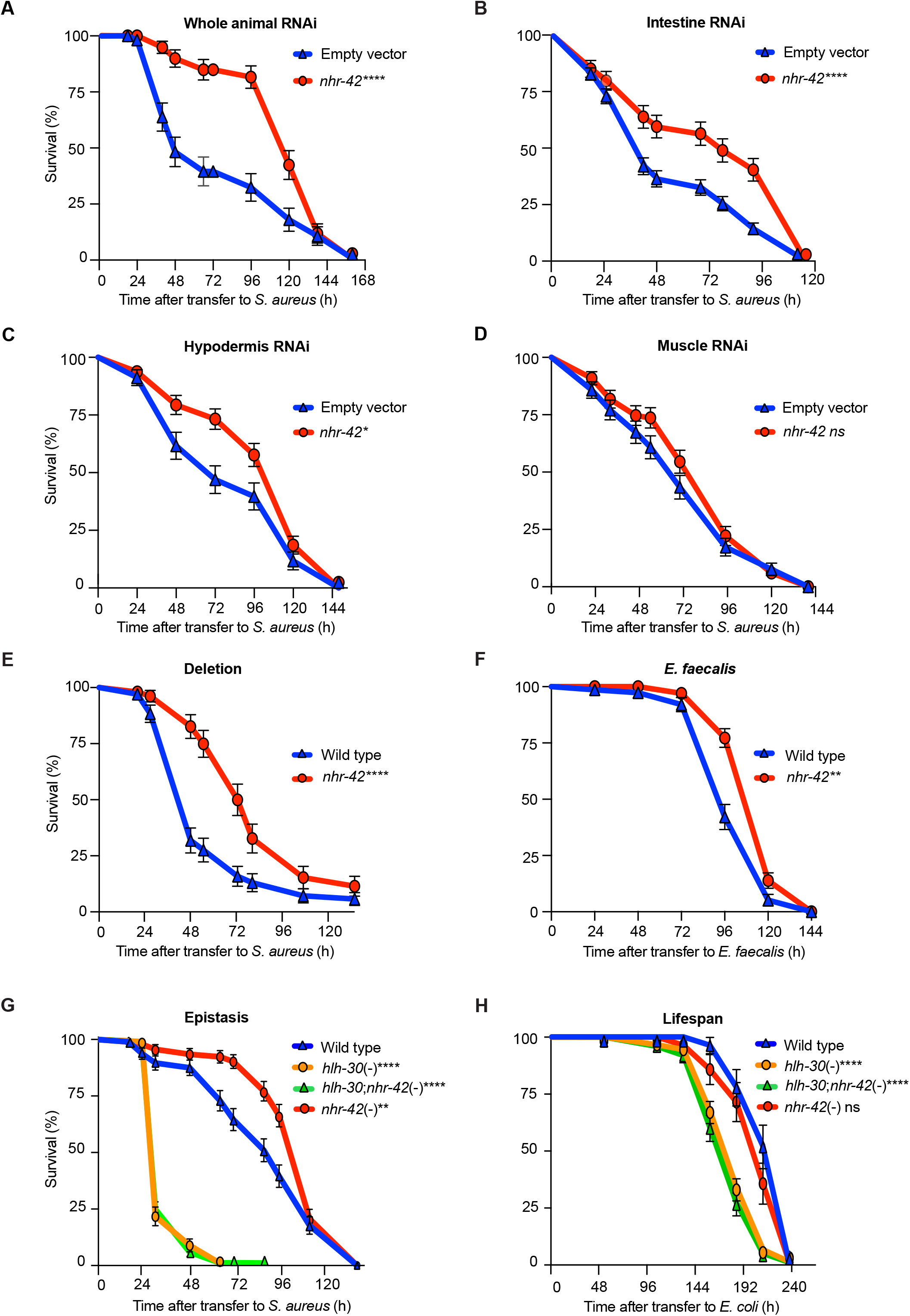
*nhr-42* represses infection resistance. **(A)**Survival of wild type animals fed *E. coli* expressing RNAi empty vector or *nhr-42* double stranded DNA upon *S. aureus* SH1000 infection. Data are representative of two independent replicates. **** *P* ≤ 0.0001. Log-Rank. **(B)**Survival of intestine specific RNAi animals MGH-171 fed *E. coli* expressing RNAi empty vector or *nhr-42* double stranded DNA upon *S. aureus* SH1000 infection. Data are representative of two independent replicates. **** *P* ≤ 0.0001. Log-Rank. **C)**Survival of hypodermis specific RNAi animals NR222 fed *E. coli* expressing RNAi empty vector or *nhr-42* double stranded DNA upon *S. aureus* SH1000 infection. Data are representative of two independent replicates. * *P* ≤ 0.05. Log-Rank. **(D)**Survival of muscle specific RNAi animals NR350 fed *E. coli* expressing RNAi empty vector or *nhr-42* double stranded DNA upon *S. aureus* SH1000 infection. Data are representative of two independent replicates. Log-Rank. **(E)**Survival of wild type and *nhr-42* deletion mutant (tm1375) upon *S. aureus* SH1000 infection. **** *P* ≤0.0001. Log-Rank. Data are representative of 3 independent replicates **(F)**Survival of wild type and *nhr-42* deletion mutant (tm1375) upon *E. faecalis* infection. ** *P* ≤ 0.01 Log-Rank. Data are representative of 3 independent replicates. **(G)**Survival of wild type, *nhr-42* loss of function mutant, *hlh-30* loss of function mutation, and *nhr-42*;hlh-30 double mutant upon *S. aureus* infection. Wild type compared to *hlh-30*(-) *** *P* ≤ 0.001. Wild type compared to *nhr-42*(-) ** *P* ≤ 0.01. Wild type compared to *hlh-30;nhr-42*(-) *** *P* ≤ 0.001. Log-Rank. Data are representative of 3 independent replicates. **(H)**Lifespan of *nhr-42* loss of function mutant, *hlh-30* loss of function mutation, and *nhr-42*;*hlh-30* double mutant fed non-pathogenic *E. coli* OP50. Wildtype compared to *hlh-30*(-) *** *P* ≤ 0.001, Wild type compared to *hlh-30*;*nhr-42*(-). Data are representative of 2 independent replicates. Log-Rank.

We tested if *nhr-42* mutants were resistant to other pathogens, such as *Pseudomonas aeruginosa* and *Enterococcus faecalis. nhr-42* mutants were not resistant to *P. aeruginosa* compared to wild type (Supplemental Figure 2a) but were more resistant to *E. faecalis* (Figure 2f). These data suggest that *nhr-42* limits defense against gram-positive pathogen infection.

*hlh-30* loss confers hypersusceptibility to *S. aureus* infection (Visvikis et al., 2014), while *nhr-42* loss of function confers the opposite. *nhr-42*;*hlh-30* double mutants were hypersusceptible to infection, similar to *hlh-30* single mutants (Figure 2g) implying that *hlh-30* is epistatic to *nhr-42*. Uninfected *nhr-42*;*hlh-30* mutants had a shorter lifespan compared to wild type animals, similar to that of *hlh-30* (Figure 2h). Importantly, *nhr-42* loss of function alone had no detrimental effects on lifespan (Figure 2h). These data are consistent with the idea that *nhr-42* represses genes that require *hlh-30* for their expression.

### *nhr-42* regulates genes involved in host defense and metabolism

Since *nhr-42* deficient animals are resistant to infection, we hypothesized that *nhr-42* may repress host defense genes. Whole animal RNA-sequencing analysis (Figure 3a) of uninfected animals showed that around 300 genes were upregulated in *nhr-42* mutants (Supplementary table 1) compared to wildtype. (Figure 3c, d). GO over-representation analysis showed enrichment of innate immunity, host defense, cuticle structure and FLP neuropeptide signaling genes/categories (Figure 3g). Seventeen (∼6%) of these genes were upregulated by HLH-30 during infection (Figure 3b) (Visvikis et al., 2014).

**Figure 3.**
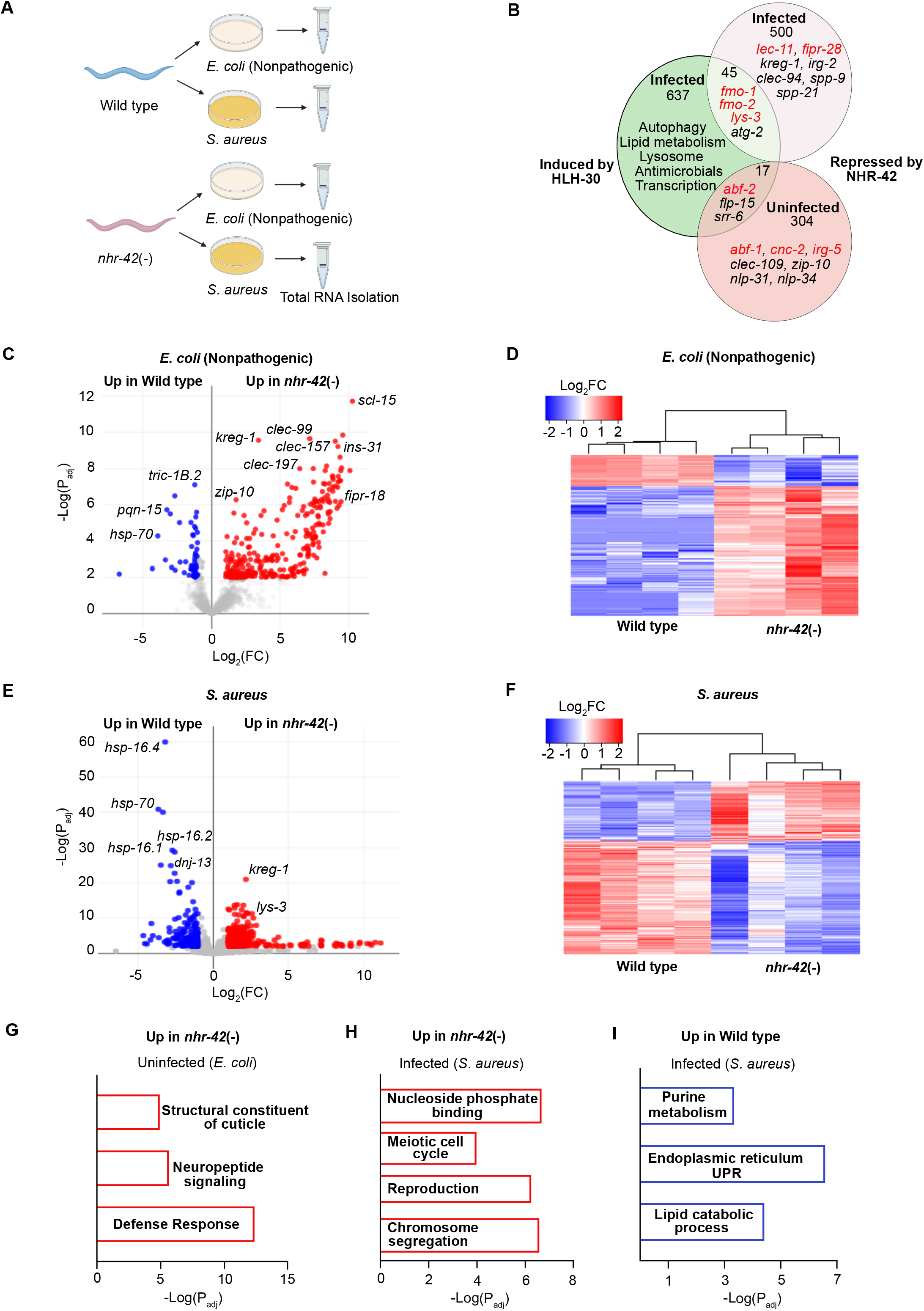
*nhr-42* regulates genes involved in host defense and metabolism. **(A)**Schematic of experimental design for RNA sequencing of wild type and *nhr-42* loss of function mutants fed *E. coli* or infected with *S. aureus* for 5 hours. **(B)**Venn diagram of genes induced by HLH-30 upon infection and genes repressed by NHR-42 under uninfected and infected conditions. Shaded area represents genes in common. **(C-D)** Volcano plot of differentially expressed genes in *nhr-42* loss of function mutants compared to wild type when fed non-pathogenic *E. coli* (*P* ≤ 0.01). Genes upregulated in *nhr-42* loss of function mutants are in red and genes upregulated in wild type are in blue. FC, fold change. P_adj_, adjusted P value. (D) Heat map of differentially expressed genes (Log_2_FC) in *nhr-42* loss of function mutants (4 biological replicates) compared to wild type (4 biological replicates) when fed non-pathogenic *E. coli*. **(E-F)** Volcano plot of differentially expressed genes in *nhr-42* loss of function mutants compared to wild type when infected with *S. aureus* (*P* ≤ 0.01). Genes upregulated in *nhr-42* loss of function mutants are in red and genes upregulated in wild type are in blue. FC, fold change. P_adj_, adjusted P value. Heat map of differentially expressed genes (Log_2_FC) in *nhr-42* loss of function mutants (4 biological replicates) compared to wild type (4 biological replicates) when infected with *S. aureus* (F). **(G)** Gene enrichment analysis of genes upregulated in *nhr-42* loss of function mutants when fed non-pathogenic *E. coli*. **(H)** Gene enrichment analysis of genes upregulated in *nhr-42* loss of function mutants when infected with *S. aureus*. **(I)** Gene enrichment analysis of genes upregulated in wild type when infected with *S. aureus*.

Our data also showed that around 600 genes were differentially regulated in infected *nhr-42* mutants compared to wild type (Figure 3e, f, supplementary Table 1). GO overrepresentation analysis identified that categories related to reproduction and cell cycle were upregulated in infected *nhr-42* mutants (Figure 3h). 45 genes upregulated in *nhr-42* mutants were previously shown to be HLH-30 regulated (Visvikis et al., 2014), including important host defense genes such as *atg-2, fmo-1, fmo-2*, and *lys-3* (Figure 3b). Conversely, categories related to endoplasmic reticulum unfolded protein response (UPR^ER^), purine, and lipid metabolism were downregulated in *nhr-42* mutants (Figure 3i).

### *nhr-42* drives lipid catabolism during infection

During infection, *C. elegans* exhibits rapid loss of neutral lipids (Dasgupta et al., 2020; Nhan et al., 2019). *nhr-42* mutants exhibited lower expression of seven lipid catabolism genes compared to wild type (Figure 4a), including predicted enoyl-CoA hydratase *ech-7* and acyl-CoA oxidases *acox1*.*3* and *acox1*.*4*, which are potentially involved in fatty acid β-oxidation. Also downregulated in *nhr-42* mutants was ABhydrolase domain-containing homolog *abhd-3*.*2*, which is predicted to be a phospholipase involved in catabolism of medium chain fatty acids. Oil Red O (ORO) staining for lipid droplets showed defective lipid mobilization in *nhr-42* mutants compared to wild type during infection (Figure 4b-c). After 8 h of infection with *S. aureus* (Figure 4d-o) or *E. faecalis* (Supplementary figure 5a-f), *nhr-42* mutants showed significantly stronger ORO staining compared to wild type. *hlh-30* mutants and *hlh-30;nhr-42* double mutants also showed defective lipid mobilization similar to *nhr-42* single mutants (Supplementary figure 4a-m). Taken together, these data suggest that *nhr-42* may mediate lipid mobilization by HLH-30 during *S. aureus* and *E. faecalis* infection.

**Figure 4.**
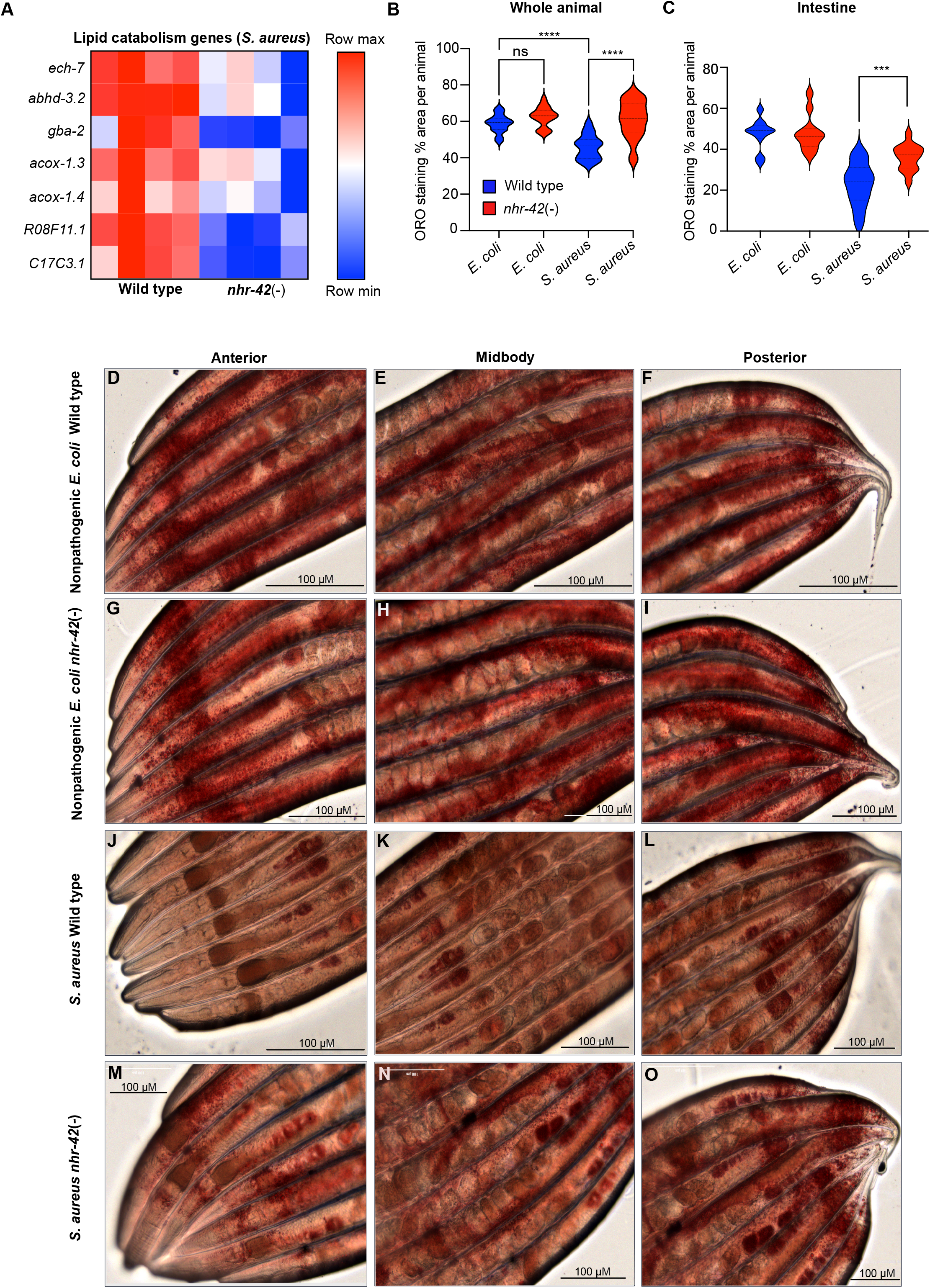
NHR-42 represses host defense genes required for enhanced survival during infection. **(A)** Heat map of host defense genes downregulated in *nhr-42* loss of function mutants compared to wild type when fed non-pathogenic *E. coli* (Uninfected) (*P* ≤ 0.01). Row normalized FPKM values. **(J-K)** RT-qPCR of mRNA transcripts of *abf-2, cnc-2, irg-5, pals-23, cnc-4*, and *lec-11* on wild type and *nhr-42* loss of function animals fed non-pathogenic *E. coli* (Uninfected). 5-8 biological replicates. n=∼1000 animals per replicate. **(B)** * *P* ≤ 0.05, ** *P* ≤ 0.01. *t*-test. **(C)**Survival of wild type and *nhr-42* loss of function mutant animals fed *E. coli* expressing RNAi empty vector or *abf-2* double stranded DNA upon *S. aureus* SH1000 infection. Data are representative of three independent replicates. Wild type EV compared to *nhr-42*(-) EV ** *P* ≤ 0.01, *nhr-42*(-) EV compared to *nhr-42*(-) *abf-2* RNAi ** *P* ≤ 0.01. Log-rank. **(D)**Survival of wild type and *nhr-42* loss of function mutant animals fed *E. coli* expressing RNAi empty vector or *cnc-2* double stranded DNA upon *S. aureus* SH1000 infection. Data are representative of two independent replicates. Wild type EV compared to *nhr-42*(-) EV *** *P* ≤ 0.001, *nhr-42*(-) EV compared to nhr-42(-) *cnc-2* RNAi *** *P* ≤ 0.001. Log-rank. **(E)**Survival of wild type and *nhr-42* loss of function mutant animals fed *E. coli* expressing RNAi empty vector or *irg-5* double stranded DNA upon *S. aureus* SH1000 infection. Data are representative of two independent replicates. Wild type EV compared to *nhr-42*(-) EV **** *P* ≤ 0.0001, *nhr-42*(-) EV compared to *nhr-42*(-) *irg-5* RNAi ns. Log-rank. **(F)**Survival of wild type and *nhr-42* loss of function mutant animals fed *E. coli* expressing RNAi empty vector or *pals-23* double stranded DNA upon *S. aureus* SH1000 infection. Data are representative of two independent replicates. Wild type EV compared to *nhr-42*(-) EV *** *P* ≤ 0.001, *nhr-42*(-) EV compared to *nhr-42*(-) *pals-23* RNAi ns. Log-rank. **(G)**Survival of wild type and *nhr-42* loss of function mutant animals fed *E. coli* expressing RNAi empty vector or *cnc-4* double stranded DNA upon *S. aureus* SH1000 infection. Data are representative of two independent replicates. Wild type EV compared to *nhr-42*(-) EV *** *P* ≤ 0.001, *nhr-42*(-) EV compared to *nhr-42*(-) *cnc-4* RNAi ns. Log-rank. **(H)**Survival of wild type and *nhr-42* loss of function mutant animals fed *E. coli* expressing RNAi empty vector or *lec-11* double stranded DNA upon *S. aureus* SH1000 infection. Data are representative of two independent replicates. Wild type EV compared to *nhr-42*(-) EV *** *P* ≤ 0.001, *nhr-42*(-) EV compared to *nhr-42*(-) *lec-11* RNAi *** *P* ≤ 0.001. Log-rank.

### *nhr-42* represses host defense genes required for enhanced survival during infection

Host defense genes upregulated in uninfected *nhr-42* mutants include antibacterial factor-related genes *abf-1* and *abf-2*, antimicrobial caenacins *cnc-2, cnc-4, and cnc-6*, immune response gene *irg-5*, as well as *lec-11* and *pals-23*, which were previously implicated in host defense (Zehrbach et al., 2017)(Kato et al., 2002) (Nandakumar and Tan, 2008) (Bui and Troemel, 2020) (Figure 5a). RT-qPCR independently confirmed these results except for *irg-5* (Figure 5b). Knockdown of host defense genes *irg-5, pals-23, cnc-4*, and *lec-11* had no effect on survival of *nhr-42* mutants (Figure 5e, f, g, and h). In contrast, knockdown of *cnc-2* and *abf-2* abolished their infection resistance phenotype (Figure 5c, d). This suggests that higher *abf-2* and *cnc-2* are essential for the infection resistance phenotype of *nhr-42* mutants. Together these data show that *nhr-42* is a negative regulator of host defense genes that promote survival of infection.

**Figure 5.**
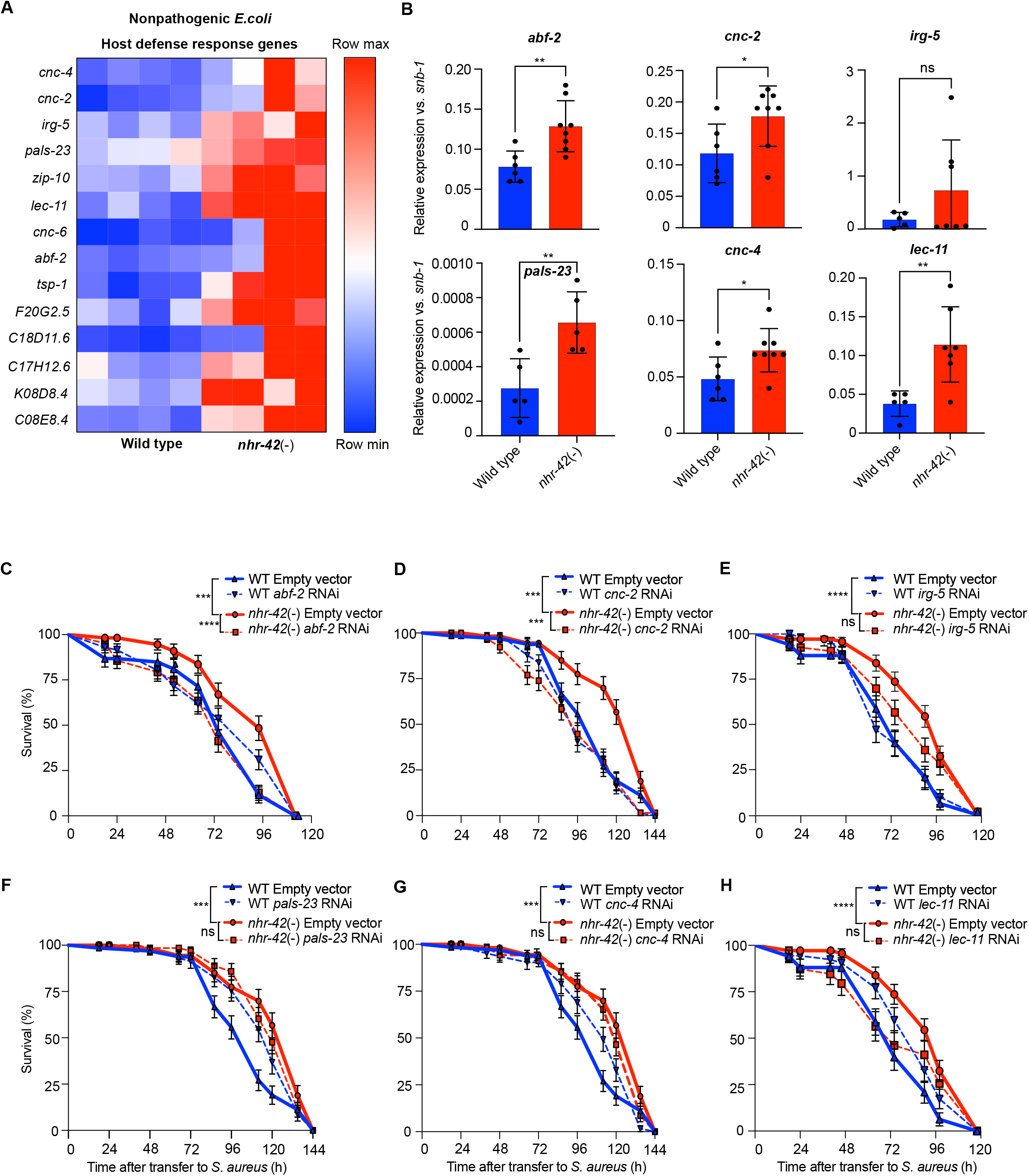
*abf-2* mediates infection resistance in *nhr-42* mutants. **(A)**Colony forming unit (CFU) counts of wild type and *nhr-42* loss of function mutants upon *S. aureus* infection, 6,12, and 24 h. ** *P* ≤ 0.01. *t*-test. **(B)**CFU count of wild type and *nhr-42* loss of function mutant animals fed *E. coli* expressing RNAi empty vector or *abf-2* upon *S. aureus* infection 24h. * *P* ≤ 0.05, ** *P* ≤ 0.01. *t*-test. **(C)**RT-qPCR of *abf-2* transcript in wild type, *nhr-42* loss of function mutant, *hlh-30* loss of function mutant, and *nhr-42*;*hlh-30* loss of function mutants. ns. *t*-test. **(D)**Pumping rate of *nhr-42* loss of function mutant compared to wild type. NS. *t*-test.

### *abf-2* mediates infection resistance in *nhr-42* mutants

Whether *C. elegans* antimicrobial effectors inhibit *S. aureus* growth *in vivo* is unknown. After 6 and 12 h of infection the *S. aureus* burden trended lower in *nhr-42* mutants compared to wild type and was significantly lower after 24 h (Figure 6a). Pharyngeal pumping was not significantly different between wild type and *nhr-42* mutants, ruling out decreased feeding as a source of this difference (Figure 6d). This suggested that *nhr-42* mutants resist *S. aureus* infection better than wild type. Knockdown of *abf-2* suppressed the decreased colonization phenotype of *nhr-42* mutants suggesting that *abf-2* limits infection in *nhr-42* mutants. (Figure 6b). Although *abf-2* expression was increased in *nhr-42* mutants prior to infection, *abf-2* expression levels remained unchanged during infection (Figure 6c). These data suggest that enhanced constitutive expression of *abf-2* prior to infection is essential for limiting pathogen burden in *nhr-42* mutants.

**Figure 6.**
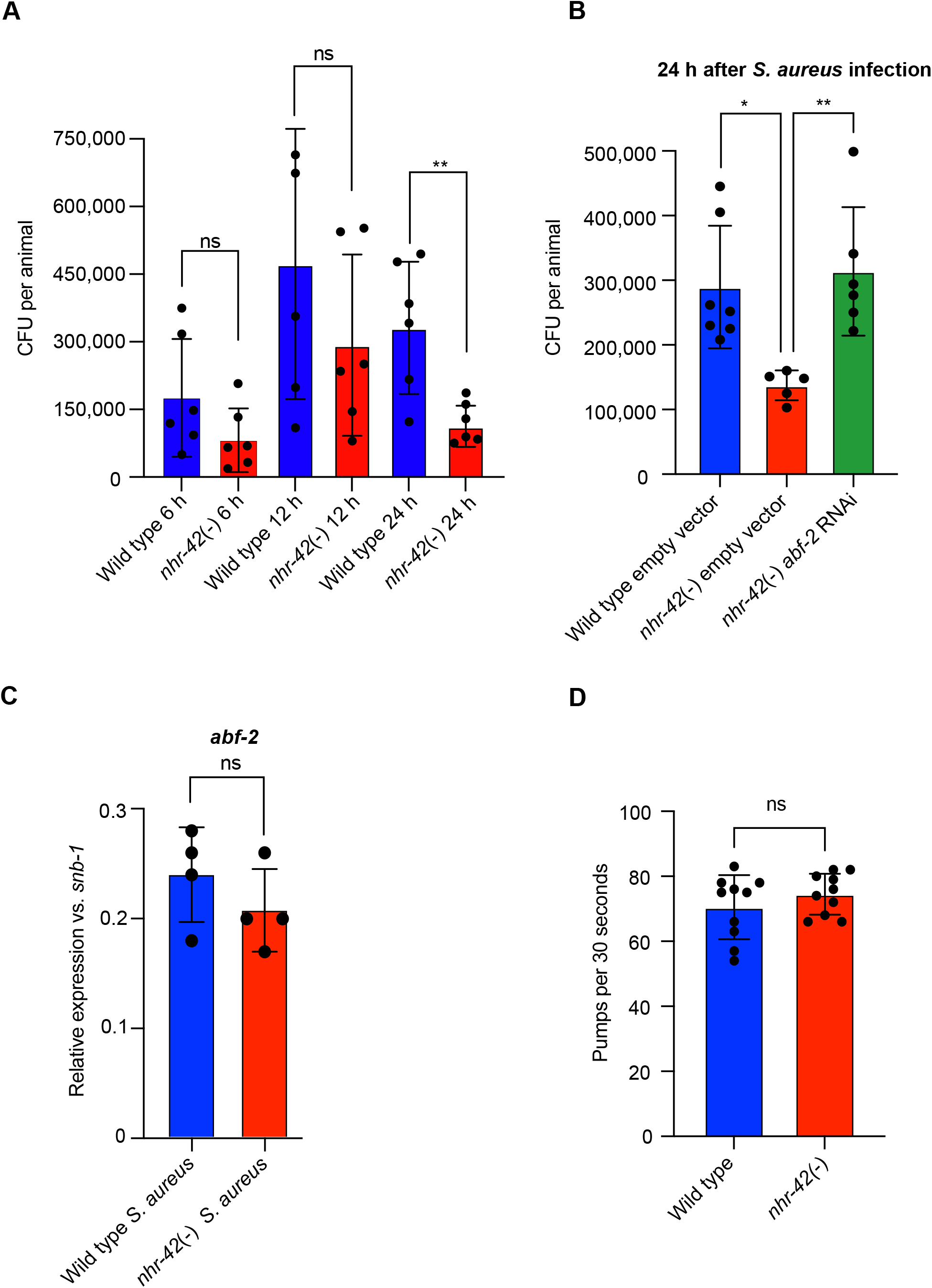
*nhr-42* drives lipid catabolism during infection. **(A)**Heat map of lipid catabolism genes upregulated in *nhr-42* loss of function mutants compared to wild type when fed non-pathogenic *E. coli* (Uninfected) (P ≤ 0.01). Row normalized FPKM values. **(B)**Quantification of Oil Red O staining in wild type and *nhr-42* loss of function animals. (D-O). **** *P* ≤ 0.0001. One-way ANOVA. Representative of 3 independent replicates, 10-20 animals per replicate. **(C)**Quantification of Oil Red O staining in intestinal tissue of wild type and *nhr-42* loss of function animals. *** *P* ≤ 0.001. One-way ANOVA. Representative of 3 independent replicates, 10-20 animals per replicate. **(D-F)** Images of wild type animals stained with Oil Red O, fed non-pathogenic *E. coli* 10 h. Scale bar-100 micrometers. **(G-I)** Images of *nhr-42* loss of function animals stained with Oil Red O, fed non-pathogenic *E. coli*, 10 h. Scale bar-100 micrometers. **(G-I)** Images of wild type animals stained with Oil Red O, infected with *S. aureus*. 10 h. Scale bar-100 micrometers. **(J-L)** Images of *nhr-42* loss of function animals stained with Oil Red O, infected with *S. aureus*. 10 h. Scale bar-100 micrometers.

## Discussion

This study identifies novel nuclear receptor *nhr-42* to be a negative regulator of host defense and a positive regulator of lipid catabolism, which functions downstream of *hlh-30*. We find that *nhr-42* plays at least two important roles: 1. In non-infected animals, *nhr-42* downregulates host defense genes required for resistance to infection, and 2. during infection *hlh-30* dependent induction of *nhr-42* causes upregulation of genes involved in lipid catabolism, driving lipid mobilization. (Figure 7).

**Figure 7.**
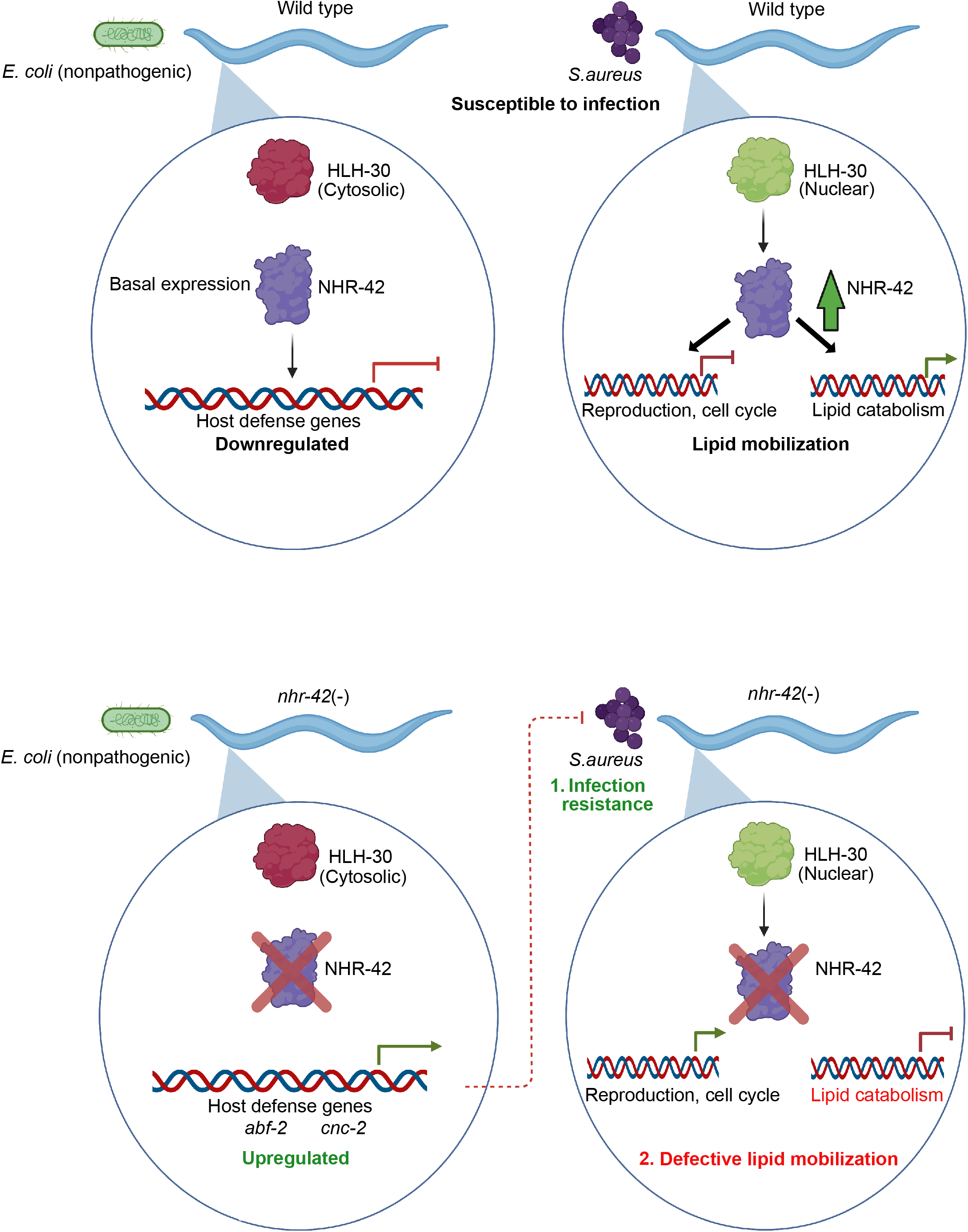
Working model. During infection, induction of *nhr-42* expression by *hlh-30* leads to upregulation of several gene categories, including lipid catabolism which leads to lipid mobilization. Conversely, *nhr-42* downregulates genes involved in reproduction and cell cycle during infection. In *nhr-42* mutants: 1. host defense genes such as *abf-2* and *cnc-2* are upregulated constitutively, making these animals more resistant to infection, 2. after infection, lower expression of genes involved in lipid catabolism possibly leads to defective lipid mobilization.

Several transcription factors, including HLH-30, ATF-7 and NHR-86, have been previously identified, which upregulate host defense genes upon infection (Fletcher et al., 2019; Peterson et al., 2019; Visvikis et al., 2014), but negative regulators of host defense genes are poorly understood in *C. elegans*. Our study identifies *nhr-42* as the first transcription factor that represses several genes involved in innate immunity.Previous studies have shown that several of these repressed genes are induced upon infection and are key in promoting host survival of infection. For example, overexpression of the antimicrobial peptide *cnc-2* during *Drechmeria coniospora* infection enables animals to survive better compared to wild type (Zehrbach et al., 2017). Furthermore, *abf-2* and *lec-11* have been shown to kill bacteria *in vitro* (Kato et al., 2002) (Nandakumar and Tan, 2008). *abf-2*, an ASABF-type antimicrobial peptide is expressed in the pharynx, similar to *nhr-42* and is predicted to be secreted into the intestine (Kato et al., 2002). *abf-2* is induced by *S. aureus, P. aeruginosa*, and *Salmonella typhimurium* infection (Kato et al., 2002). *abf-2* knockdown leads to higher burden and persistence of *S. typhimurium* infection compared to wild type animals (Alegado and Tan, 2008). Moreover, recombinant *abf-2* is bactericidal against *S. aureus* and several other bacterial pathogens *in vitro* (Kato et al., 2002). We also saw upregulation of genes involved in maintaining the structural integrity of the collagenous cuticle in *nhr-42* mutants (Figure 3g). Several collagen genes were shown to be required for enhanced survival during *P. aeruginosa* infection (Sellegounder et al., 2019).

Our data suggest that the upregulation of such antimicrobials in *nhr-42* mutants, results in enhanced survival and lower pathogen burden after infection. Interestingly, *abf-2* expression is induced upon infection in an *hlh-30* dependent manner, however *hlh-30* mutants have similar pathogen burden as wild type animals (Visvikis et al., 2014). Therefore, it is likely that constitutive induction of *abf-2*, and not necessarily its induction by *hlh-30*, is a key protective mechanism in *nhr-42* mutants.

We also found that other host defense genes induced by HLH-30, such as *fmo-2* were upregulated in *nhr-42* mutants (Figure 3b, supplementary Figure 6a,b). *fmo-2* is one of the most highly induced genes upon *S. aureus* infection (Wani et al., 2021). We recently showed that animals lacking *fmo-2* are susceptible to *S. aureus* infection and that overexpression of *fmo-2* is protective (Wani et al., 2021). Together these data suggest that *nhr-42* mutants are primed for host defense response even when not infected. A combination of higher expression of host defense genes in the absence of infection and during infection likely enables *nhr-42* deficient animals to resist infection and enhance survival.

Recent studies have highlighted the importance of certain fatty acids, such as oleate, in regulating host defense responses in *C. elegans* (Anderson et al., 2019). Importantly, *nhr-42* upregulation by *hlh-30* during infection leads to higher expression of hypothetical lipid catabolism genes potentially involved in β -oxidation. Lipid droplet staining showed that *nhr-42* mutants are defective in lipid mobilization during infection. Increased lipid storage as a result of a high glucose diet has been shown to help *C. elegans* survive *E. faecalis* infection (Dasgupta et al., 2020). In contrast, somatic lipid mobilization during *P. aeruginosa* infection is required for the host defense response (Nhan et al., 2019).However, *nhr-42* mutants are resistant to infection despite having similar defects in lipid mobilization to *hlh-30* mutants and *hlh-30;nhr-42* double mutants, which are hypersusceptible to infection. This suggests that the infection resistance seen in *nhr-42* deficient animals maybe independent of lipid mobilization and supports the idea that *nhr-42* may mediate lipid mobilization downstream of *hlh-30*.

Oxidative and nutrient stress drive lipid mobilization mediated by SKN-1/NRF2 and HLH-30/TFEB respectively (Lynn et al., 2015; O’Rourke and Ruvkun, 2013).Interestingly, it was recently shown that *P. aeruginosa* infection induced lipid mobilization is also driven by SKN-1/NRF2 (Nhan et al., 2019). The same study found that *nhr-42* is induced in *skn-1* gain of function animals, which are prone to accelerated somatic lipid mobilization during infection. This suggests that *nhr-42* could function as a driver of lipid mobilization during various infections. Lipid mobilization during stress could serve several purposes (Hou and Taubert, 2012a). Lipids can act as signaling molecules or structural elements of cellular membranes, and influence energy metabolism and reproduction to better adapt to stress. Consistently, several lipid metabolism enzymes and factors that drive lipid mobilization have been linked to extended lifespan in *C. elegans* (Hou and Taubert, 2012b). The physiological roles of lipid metabolism during *S. aureus* infection deserve further study.

*nhr-42* could be important for limiting the host defense response after infection. In the wild, *C. elegans* can behaviorally avoid pathogenic microbes and, therefore, mechanisms that downregulate the host response may be beneficial to reproduction or survival. *nhr-42* expression peaks late during infection and thus *nhr-42* could work in a negative feedback loop to downregulate genes such as *abf-2, fmo-2 and lys-3* that are induced upon infection by *hlh-30* (Figure 3b). Negative regulation of innate immunity in *C. elegans* remains poorly understood. Our study identifies and characterizes a previously unknown orphan nuclear receptor *nhr-42*, as a negative regulator of host defense genes. Our data also highlights how *hlh-30* is able to indirectly regulate gene sets involved in host defense and lipid catabolism through *nhr-42* after infection. It will be important to determine if nuclear receptors downstream of TFEB also regulate innate immunity and metabolism during bacterial infection in mammals.

## Acknowledgements

The authors are grateful to the members of the Irazoqui laboratory and Department of Microbiology and Physiological Systems for helpful insights and discussions. Authors would like to thank Michael Francis and Craig Mello laboratory members-Christopher Lambert, Krishna Ghanta, and Daniel Durning for their help in providing and generating reagents for this study. Authors are also grateful to Alper Kucukural for his help with RNA-sequencing data analysis.

## Author contributions

D.G. conceived and designed the experiments. D.G. analyzed the data. D.G and S.A.L. performed the experiments. All authors participated in manuscript writing and editing.

## Methods

### *C. elegans* strains and growth

All strains used in this study are detailed in table 1. *C. elegans* was grown on nematode-growth media (NGM) plates seeded with *E. coli* OP50 at 20°C according to standard procedures (Brenner, n.d.), unless mentioned otherwise.

**Table 1.**
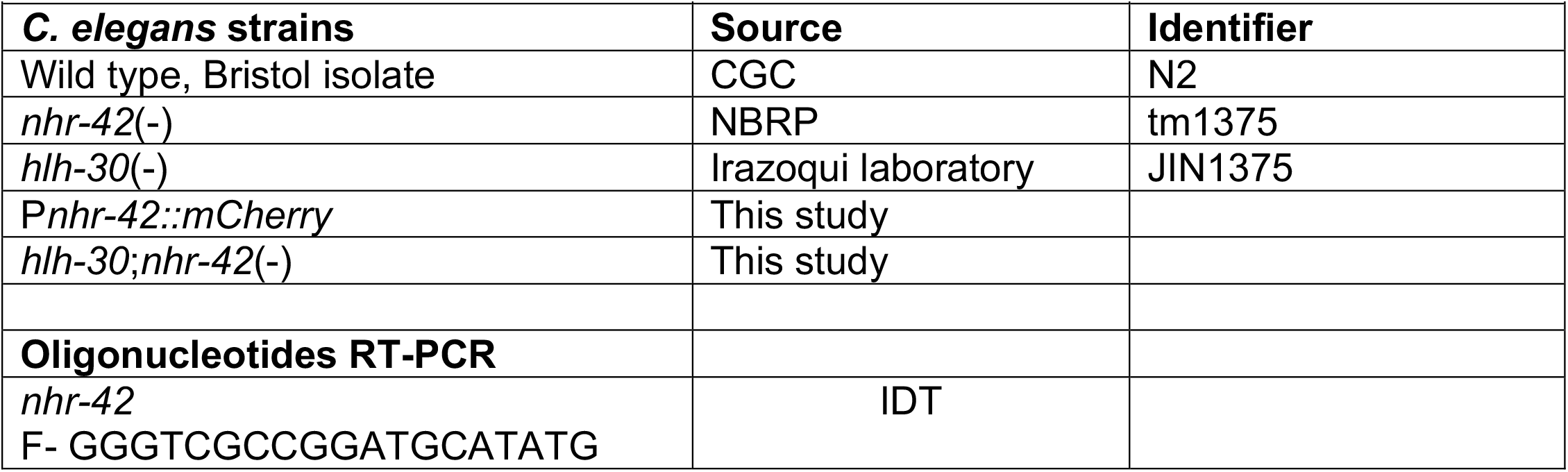
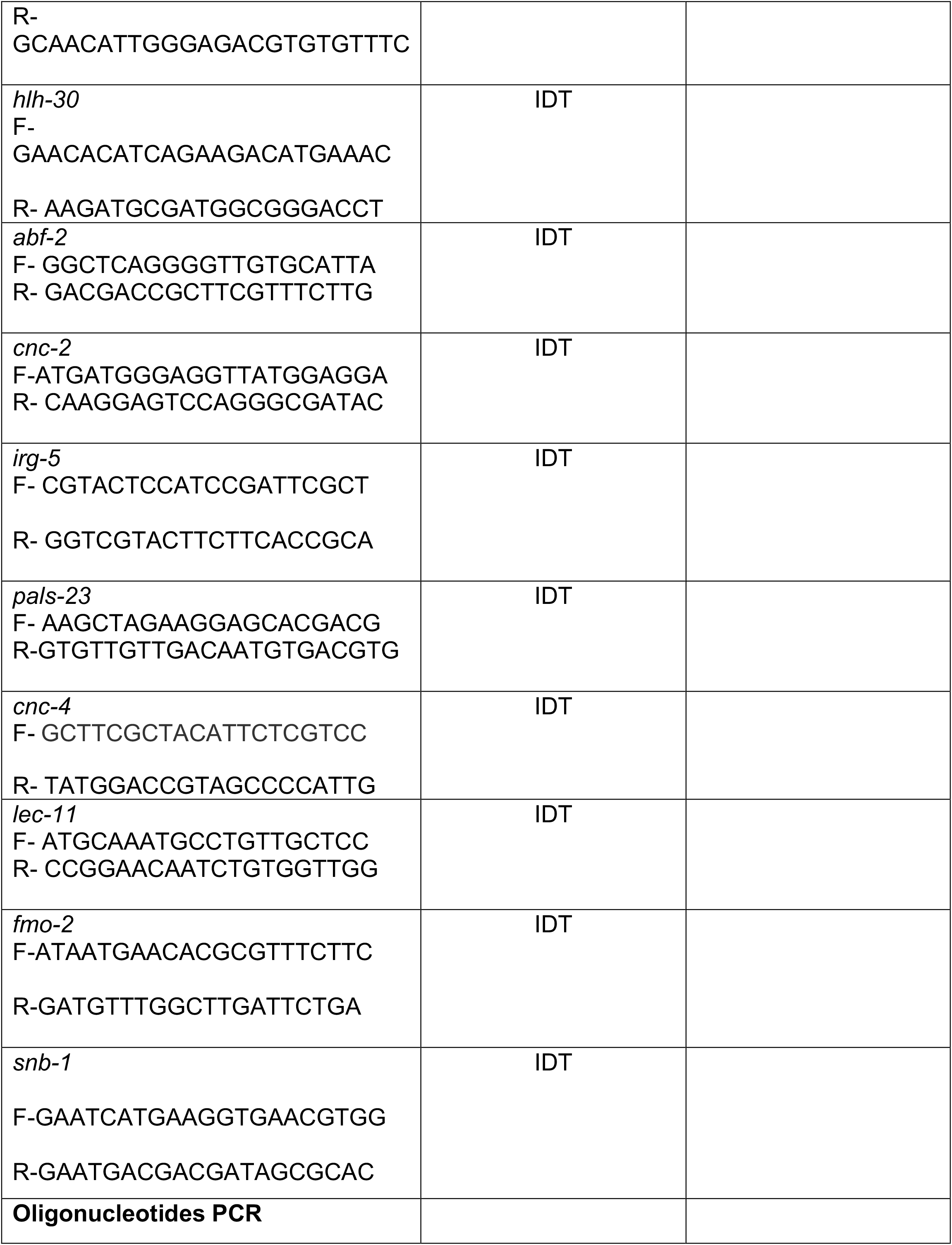
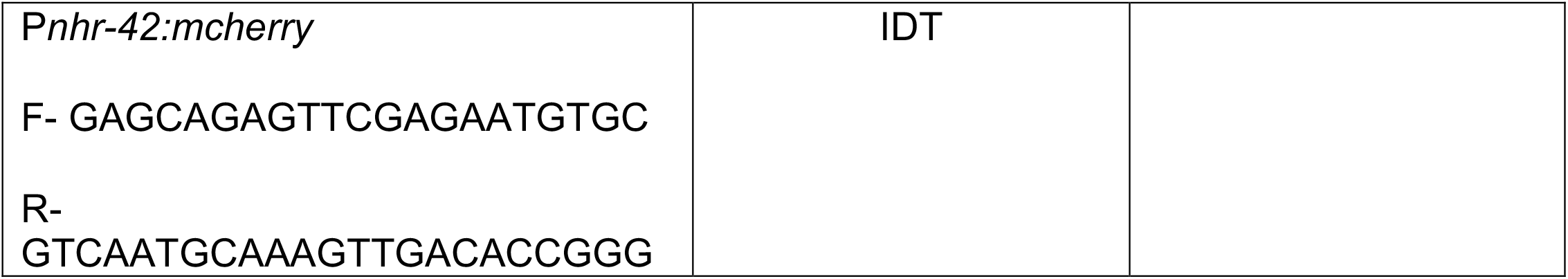

### *C. elegans* bacterial infection assays

#### *Staphylococcus aureus* infection

A single colony of SH1000 strain was grown in tryptic soy broth (TSB) supplemented with 50 µg/ml kanamycin overnight at 37°C shaking at 200-220 RPM. 10 µl of the overnight culture was spread on tryptic soy agar (TSA plates) supplemented with 10 ug/ml kanamycin. These plates were grown at 37°C for 5 h and then shifted to 25°C overnight. *C. elegans* were treated with 100 μg/ml 5-fluoro-2′-deoxyuridine (FUDR) at L4 larval stage overnight at 15 °C before transfer to *S. aureus* TSA plates with full lawn of bacteria. Three TSA plates each containing 30-40 animals were examined for each genotype of *C. elegans*. Infection assays were carried out at 25°C. For RNAi experiments animals were grown to L4 larval stage on *E. coli* strain HT115 expressing double-stranded RNA for the target genes at 20°C, treated with 100 μg/ml FUDR and transferred to 15°C overnight and subsequently used for infection assays. Survival was quantified using standard methods (Wollenberg et al., 2013). Animals dying of disrupted vulva or crawling on walls were censored for analysis. Infection assays were performed 2-3 times.

#### *Pseudomonas aeruginosa* infection

Slow killing plates were made using single colony of PA14 strain was grown overnight in LB broth 10 μl of overnight culture was spread onto slow killing assay plates and incubated at 37°C for 24 h. Plates were then placed at 25°C for a day to obtain a full bacterial lawn. Animals were grown to L4 stage on OP50 in NGM plates at 20°C. 100 μg/ml FUDR was added at L4 stage. 30-40 L4 animals of all genotypes were transferred to infection plates. 200 μg/ml FUDR was added to the infection plates. Animals were scored according to established protocols (Tan et al., 1999). Animals dying of disrupted vulva or crawling on walls were censored. Infection assays were performed 3 times.

#### *Enterococcus faecalis* infection. A

Single colony of V583 strain of *E. faecalis* was grown in BHI broth at 37°C for 6-8 h. 10 μl of this culture was spread onto BHI agar plates containing 10 μg/ml kanamycin and incubated overnight at 37°C (Yuen and Ausubel, 2018). 30-40 L4 animals grown at 20°C were then transferred onto infection plates. Animals dying of disrupted vulva or crawling on walls were censored for analysis. Infection assays were performed 3 times.

#### Longevity assays

*C. elegans* was grown at 20°C until L4 stage on NGM plates seeded with OP50 and then 100 μg/ml of FUDR was added. Animals were transferred to 25°C. Dead animals were counted every 24-48 hours. Animals that did not respond to touch with a pick were considered dead. 2-3 replicates with 20-30 animals on each plate was used for each experiment. Animals that died due to unnatural causes such as disrupted vulva or crawling on walls were censored for analysis.

#### RT-qPCR

Total RNA isolation was performed using Pure link RNA kit (Invitrogen). Animals were washed twice using sterile water and collected in 1.5 ml tubes containing 1000 μl TRIzol (Sigma-aldrich) for lysis. 100 μl chloroform was added and centrifuged at 12,000 rpm to separate aqueous phase containing RNA. The aqueous phase was transferred to Pure link RNA kit columns and purified total RNA was obtained following manufacturer’s instructions. cDNA was made using iScript cDNA synthesis kit (Bio-Rad). RT-qPCR was performed using SYBR Green Super mix (Bio-Rad) using a ViiA7 Real-Time qPCR system (Applied Biosystems). Test genes in *C. elegans*, were normalized to *snb-1*. Fold change was calculated using the Pfaffl method.

#### RNA sequencing

Total RNA was collected from *C. elegans* using Pure link RNA kit as described above. Total RNA from 4 biological replicates for each condition was sent to BGI Genomics for library preparation and sequencing using BGI-seq 500. BGI provided clean reads in FASTQ format. FASTQ files were analyzed by DolphinNext high throughput sequence analysis software (Yukselen et al., 2020). Steps followed in DolphinNext are as follows: FastQC was used to create quality control outputs. FASTQ files were aligned to *C. elegans* genome using STAR. STAR is used to count or filter out common RNAs (eg. rRNA, miRNA, tRNA, piRNA). RSEM was used to align reads to reference transcripts and estimate gene and isoform expression levels. Genome-wide Bam analysis was done by RseQC. DEBrowser v1.22.5 was used for batch effect correction and normalization. Data analysis including generation of volcano plots and heat maps was done using DE analysis within DEBrowser v1.22.5. Adjusted p-value of 0.01 was considered significant for differential gene expression. Gene Ontology representation analysis was performed using the enrichment analysis tool in Wormbase (Angeles-Albores et al., 2016) and g:profiler software (Reimand et al., 2007).

### Construction of strains

To generate P*nhr-42::mCherry*, the *nhr-42* promoter region (2000 bp upstream of the transcription start site) was amplified and cloned into entry vector pTOPO for gateway cloning. This amplified region was inserted into destination vector pDEST16 expressing *mCherry* to generate the expression vector. The expression vector was sequenced to verify proper insertion of *nhr-42* promoter region upstream of mCherry. Microinjections were carried out following standard procedures. (Dokshin et al., 2018)

### Image acquisition

Images were taken using a Lionheart FX automatic microscope (BioTek Instruments) at 4x and 20x magnification. Fluorescence microscopy-30-50 *C. elegans* were used for each experimental condition and 10-20 animals were anesthetized using 100 mM NaN_3_ on a 2% agar pad for imaging. All images were captured at the same exposure and intensity. Greyscale images were used for image analysis. A region of interest (ROI) was drawn around the pharynx which showed fluorescent expression of *mCherry*. Mean fluorescent intensity of the ROI was determined by using Analyze>Measure function in ImageJ. MFI values were plotted in GraphPad Prism and statistical analyses were performed. Fluorescence microscopy experiments were repeated 2 times.

### Lipid staining

*C. elegans* was grown at 20°C until L4 stage on NGM plates seeded with OP50.Thereafter, 20-30 animals were placed on TSA plates containing a full lawn of *S. aureus* or BHI plates for *E. faecalis* infection. *S. aureus* and *E. faecalis* infection plates were made as described. 8 hours after infection animals were washed off with water and collected in centrifuge tubes. Lipid staining using Oil Red O was performed as described (Escorcia et al., 2018). 10-20 animals were used for imaging. Animals were mounted on agarose pads and visualized using a Lionheart microscope at 4x and 20x magnification. The amount of Oil Red O staining was determined using ImageJ. Greyscale .tiff images were used for quantification. A relative threshold for the detection of red color staining was set. (Image>Threshold>). The set threshold was consistent for all images quantified. A region of interest (ROI) was drawn around each whole animal. The percent area stained inside the ROI by Oil Red O for each whole animal was calculated (Analyze>Measure>Area%). For intestine specific quantification the intestine was used as the ROI. Percent of area (Area%) stained by Oil Red O per animal was plotted and statistical analysis were performed.

### Bacterial colony forming units upon infection

*C. elegans* was grown at 20°C until L4 stage on NGM plates seeded with OP50 or RNAi clones as mentioned. Thereafter, 30-40 animals were transferred to TSA plates containing a full lawn of *S. aureus*. At the desired timepoint 10 alive animals were collected in centrifuge tubes containing autoclaved water. Colony forming units were determined as per (Visvikis et al., 2014). 5-7 biological replicates (10 animals per replicate) were used for each condition.

### Quantification and statistical analysis

Graph Pad Prism 9 was used for statistical analysis. Survival curves were compared using Log-Rank (Mantel-Cox test). *P* ≤ 0.05 was considered significant. Two-tailed *t*-tests was performed to compare with a single reference. *P* ≤ 0.05 was considered significant. For multiple comparisons, one-way ANOVA was used to establish significance. *P* ≤ 0.05 was considered significant.

---

**Supplemental Figure 1.**
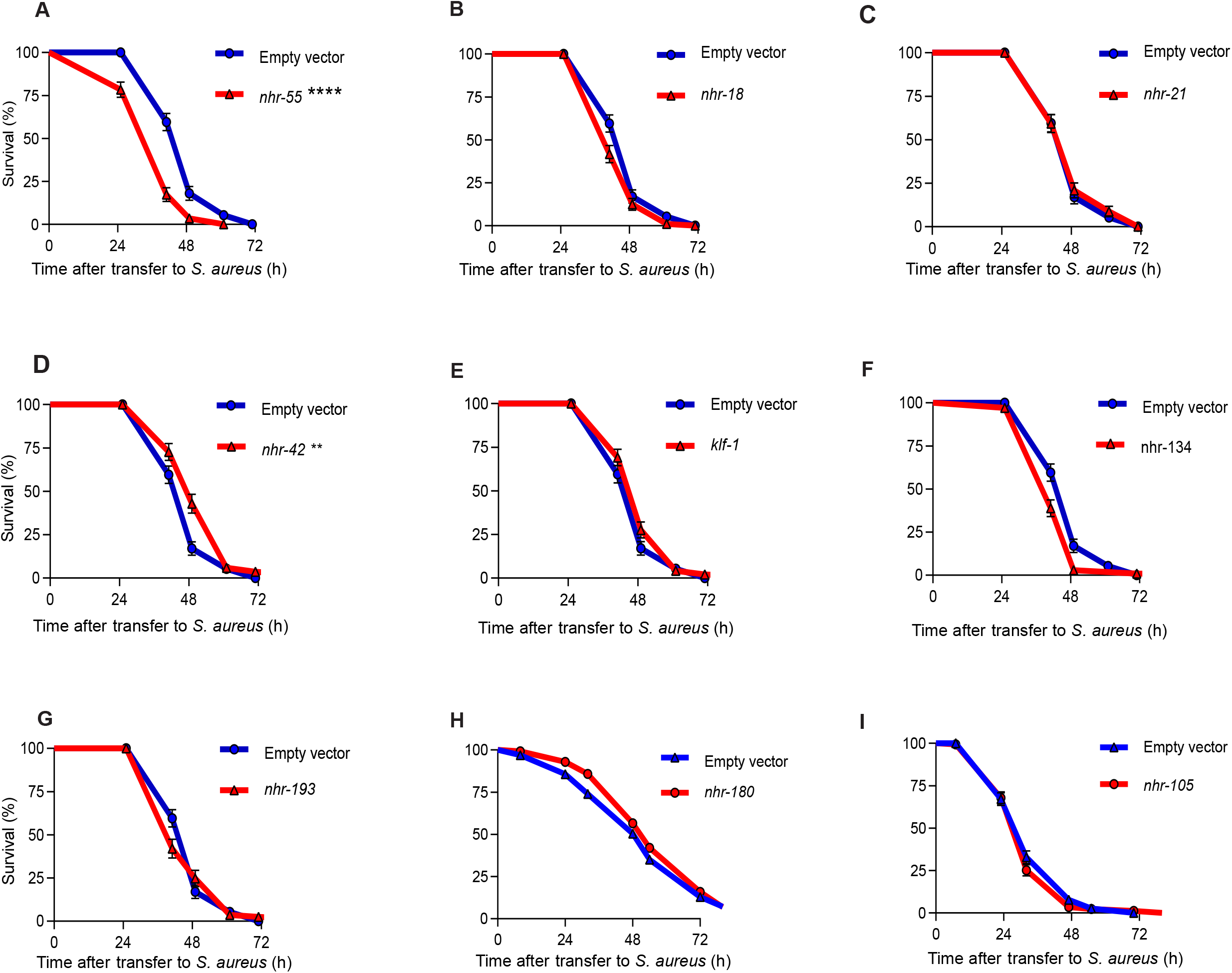
**(A-I)** Survival of wild type animals fed *E. coli* expressing RNAi empty vector or *nhr-55, nhr-18, nhr-21, nhr-42, klf-1, nhr-134, nhr-193, nhr-180, and nhr-105* double stranded DNA upon *S. aureus* SH1000 infection. ** *P* ≤ 0.01, **** *P* ≤ 0.0001. Log-Rank.

**Supplemental Figure 2.**
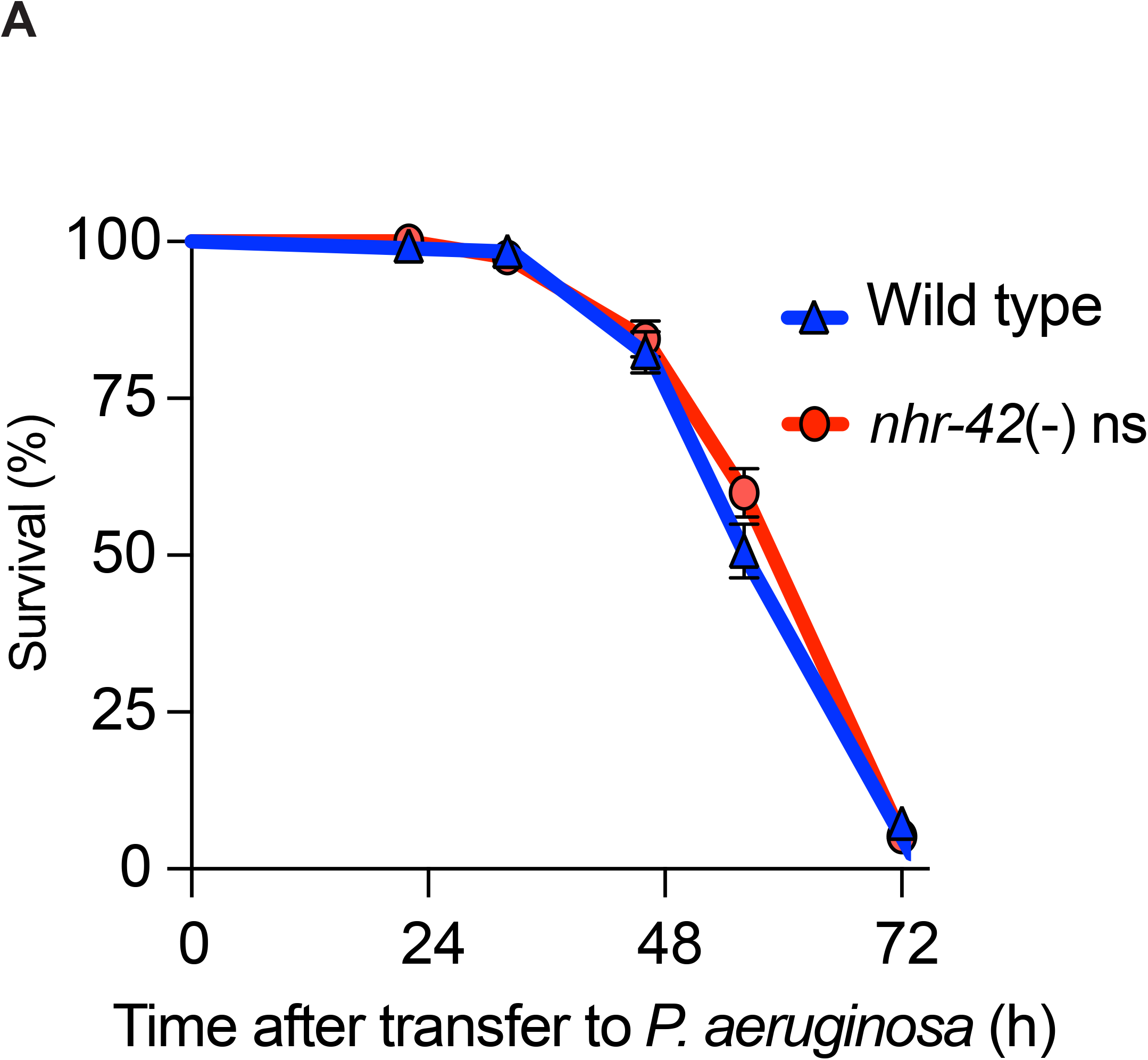
**A)**Survival of wild type and *nhr-42* deletion mutant (tm1375) upon *P. aeruginosa* infection. Data are representative of 3 independent replicates. NS. Log-Rank.

**Supplemental Figure 4.**
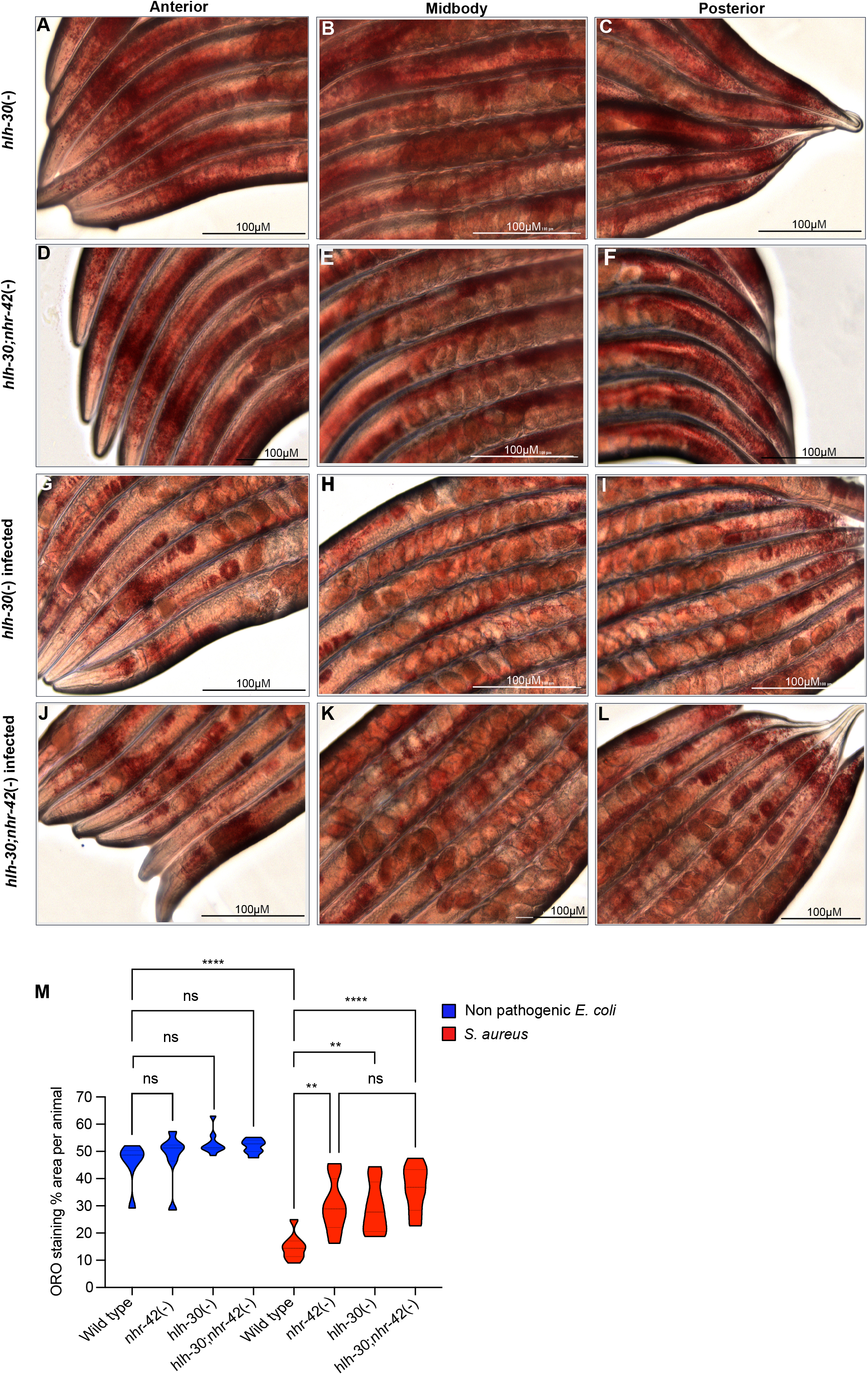
**(A-C)** Images of *hlh-30* loss of function animals stained with Oil Red O, fed non-pathogenic *E. coli* 10 h. Scale bar-100 micrometers. **(D-F)** Images of *hlh-30;nhr-42* loss of function animals stained with Oil Red O, fed non-pathogenic *E. coli*, 10 h. Scale bar-100 micrometers. **(G-I)** Images of *hlh-30* loss of function animals stained with Oil Red O, infected with *S. aureus*. 10 h. Scale bar-100 micrometers. **(J-L)** Images of *hlh-30;nhr-42* loss of function animals stained with Oil Red O, infected with *S. aureus*. 10 h. Scale bar-100 micrometers. **(M)** Quantification of images (A-O). 7-10 animals per condition. ** *P* ≤ 0.01, **** *P* ≤ 0.0001. One-way ANOVA.

**Supplemental Figure 5.**
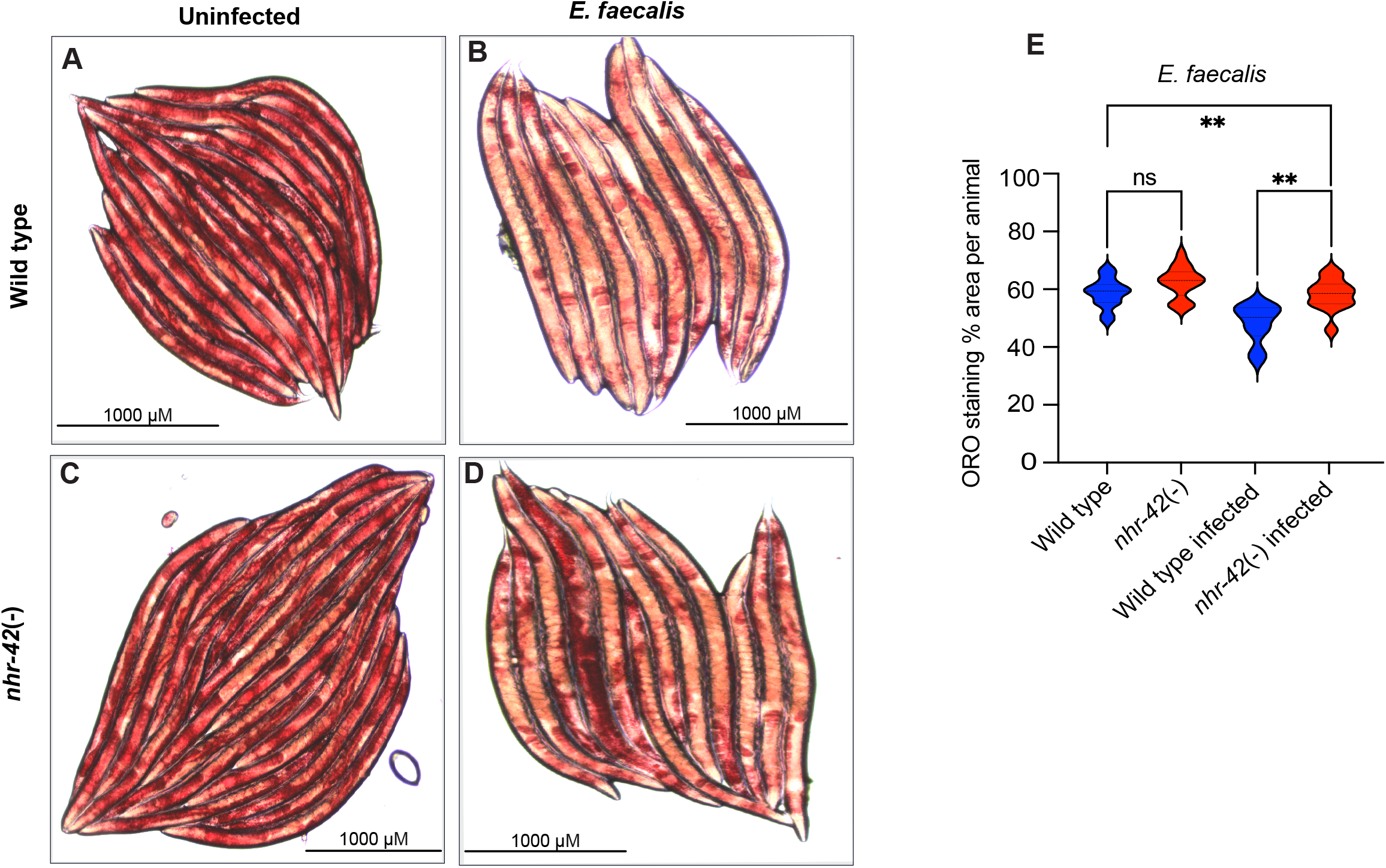
**(A)**Images of wild type animals stained with Oil Red O, fed non-pathogenic *E. coli* 10 h. Scale bar-1000 micrometer. **(B)**Images of wild type animals stained with Oil Red O, fed *E. faecalis* 10 h. Scale bar-1000 micrometer. **(C)**Images of *nhr-42* loss of function animals stained with Oil Red O, fed non-pathogenic *E. coli* 10 h. Scale bar-1000 micrometer. **(D)**Images of *nhr-42* loss of function animals stained with Oil Red O, *E. faecalis* 10 h. Scale bar-1000 micrometer. **(E)**Quantification of lipid levels in images (A-D). 10-25 animals per condition. ** *P* ≤ 0.01. One-way ANOVA.

**Supplemental Figure 6.**
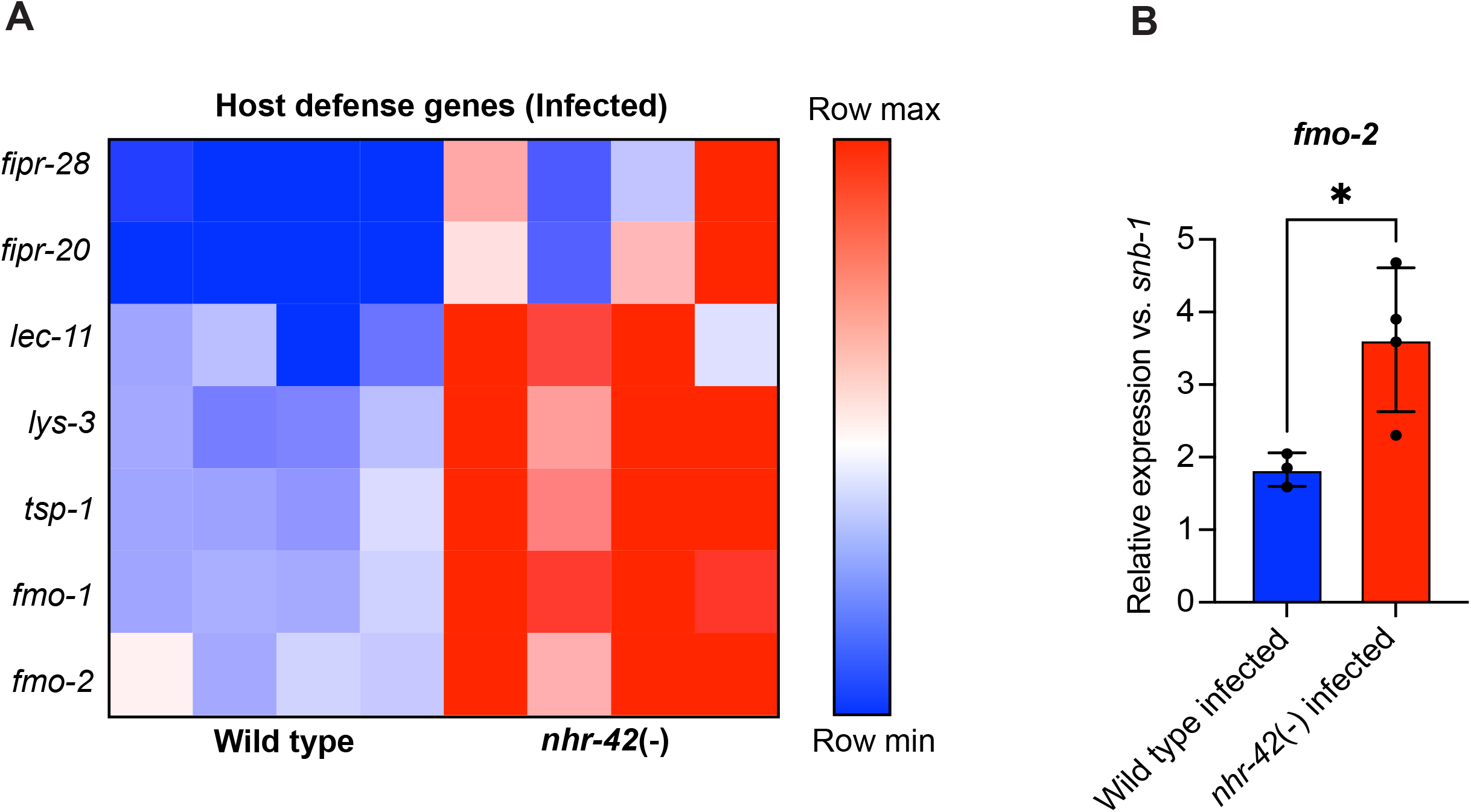
**(A)**Heat map of differentially expressed host defense genes (Log_2_FC) in *nhr-42* loss of function mutants (4 biological replicates) compared to wild type (4 biological replicates) when fed *S. aureus*. **(B)**RT-qPCR of mRNA transcripts of *fmo-2* on wild type and *nhr-42* loss of function animals infected with *S. aureus* for 5h. * *P* ≤ 0.05. *t*-test. Each dot represents a biological replicate (n=∼1000 animals)

